# Interneurons that BiTE: Harnessing the migratory capacity of cortical inhibitory interneuron precursors to treat high-grade glioma

**DOI:** 10.1101/2024.04.29.591530

**Authors:** Stephanie N. Brosius, William Manley, Kyra Harvey, Jamie Gallanaugh, Deborah Rohacek, Stewart Anderson, Thomas De Raedt

## Abstract

Glioblastoma has a poor prognosis with limited therapeutic options. To date, almost all therapeutic agents that showed promise in *in vitro* assays, preclinical trials, or even in early clinical trials have failed to make a substantial impact on the survival of patients with high-grade glioma. One major obstacle for small molecules, therapeutic proteins, or immune cells remains the blood brain barrier, which often prevents the efficient delivery of agents to the tumor. To address this delivery issue, we developed a cellular vector where implanted modified post-mitotic Migratory Cortical Inhibitory Interneuron Precursors (MCIPs) migrate to high-grade glioma by chemoattraction to locally secrete a therapeutic protein. The inhibitory interneurons of the cerebral cortex originate predominantly in the ventral/subcortical portion of the telencephalic neural tube. During fetal brain development, MCIP migration is guided over long distances by chemorepulsant and chemoattractant factors. Remarkably, several MCIP chemoattractant factors are also secreted by high-grade gliomas. Indeed, our *in vitro* and *in vivo* data show that MCIPs robustly migrate to the majority of glioblastoma cell lines evaluated. As a proof of principle, we modified the MCIPs to secrete bispecific T-cell engagers (BiTEs), linking the EGFR tumor antigen to CD3 on T-cells to create an adaptor molecule that induces an anti-tumor response of resident and supplied T-cells by bridging a tumor antigen and the T-cell receptor. We find that implanted BITE-secreting MCIPs significantly extended survival of mice injected with high-grade glioma. Therefore, we conclude that the use of MCIPs as a delivery vector for therapeutic agents could revolutionize the way we treat glioblastoma, as they allow for the local delivery of therapeutic agents in high concentrations, bypassing the need for these agents to cross the blood brain barrier while reducing the risk for systemic toxicities outside of the brain.

## Introduction

Glioblastoma (GBM) is a high-grade neoplasm of the brain with a 5% five-year overall survival^1^. High-grade gliomas (HGG) like glioblastoma represent the leading cause of cancer-related mortality in the pediatric population^2^ and are one of the most recalcitrant tumors to treat in adults as well^3^. Despite significant advances in understanding of the molecular drivers of glioblastoma, treatment options remain limited^3^.

A major challenge preventing effective treatment of glioblastoma is the delivery of therapeutic agents, due in part to the brain’s privileged space in the body^4^. Affected therapies include small molecules, therapeutic proteins, and immune cells. To circumvent these brain tumor specific issues, we established a cellular delivery system by harnessing the ability of cortical interneuron precursors to migrate through the brain and by modifying them to deliver a cytotoxic therapy. Many of the Migratory Cortical Inhibitory Interneuron Precursors (MCIPs) originate in the Medial-Ganglionic Eminence (MGE) region of the embryonic brain ^5–8^. MGE-derived MCIPs differentiate from *NKX2.1*-expressing progenitors and express *LHX6* as **<u>post-mitotic</u>** MCIPs^7^. MCIPs leave the MGE region and migrate to the cortex over the course of one week in mice and 4-6 weeks in humans^9^. This migration is guided by both chemorepulsant factors, like SEMA3A^10^, and chemoattractive factors like CXCL12, NRG1, and HGF^7^. These gradients are largely absent in the post-natal brain. Remarkably, MCIPs retain their motility when transplanted in the adult cerebral cortex of mammals^11^. Transplanted MCIPs also migrate in striatum^12^, spinal cord^13^, and cerebellum^14^. This migratory capacity, together with their ability to integrate into host circuitry, has garnered interest to use them as a cell therapy for conditions like epilepsy (clinical trial: NCT05135091). Enthusiasm has been further engendered by the successful derivation of MCIPs from human stem cells and induced Pluripotent Stem Cells (iPSCs) and their use in preclinical therapeutic transplantation studies^15,16^.

Here we demonstrate that MCIPs migrate to HGG, independent of unique tumor antigens, both *in vitro* and *in vivo*. As a proof of concept, we modified MCIPs to deliver a Bispecific T-cell Engager (BiTE) targeting the tumor, thereby prolonging survival in orthotopic xenograft models of HGG.

## Results

### High-grade gliomas secrete factors that drive chemoattraction

We hypothesized that proteins secreted by HGGs would generate a gradient capable of attracting migratory cells in an antigen independent manner. To evaluate this hypothesis, we analyzed publicly available RNA expression data of 133 pediatric HGG (pHGG) (CBTN, pedcbioportal.org), and 255 normal cortex samples (GTex, gtexportal.org) to generate a list of glioblastoma secreted proteins involved in migration (expression >3x median expression in normal cortex in at least 20% of HGG samples). This analysis revealed a total of 38 proteins, including CCL2 (6^th^ highest, 23.8x expression) and MIF (2^nd^ highest, 61.3x expression) but also IGF2 and several IGFBPs (Table 1). Several of the identified chemoattractive factors were ligands of CXCR4, namely CXCL12, MIF and HMGB1. CXCR4 induces robust migration of several cell types within the brain including macrophages, microglia, and MCIPs^17^. Given that macrophages and microglia also contribute to HGG pathology, we chose to focus our attention on MCIPs.

**Table 1:** Expression analysis of genes involved in migration secreted by pediatric high-grade glioma.

| Gene | fold change over median cortex | Median expression cortex (TPM) | median expression pHGG (TPM) | 80th percentile expression pHGG |
| --- | --- | --- | --- | --- |
| CHI3L2 | 86.4 | 0.4 | 7.8 | 38.5 |
| MIF | 61.3 | 5.6 | 187.1 | 343.7 |
| IGF2 | 44.9 | 2.2 | 7.0 | 99.8 |
| SEMA3E | 25.5 | 0.5 | 2.2 | 11.5 |
| CCL2 | 23.8 | 1.5 | 9.7 | 35.4 |
| SPP1 | 23.0 | 43.9 | 264.1 | 1007.3 |
| IGFBP2 | 21.9 | 14.3 | 117.7 | 314.0 |
| C1QL4 | 21.6 | 0.3 | 0.8 | 5.8 |
| THBS1 | 20.7 | 1.3 | 8.4 | 25.9 |
| MDK | 18.6 | 8.8 | 66.1 | 163.3 |
| NRG1 | 17.1 | 0.8 | 1.9 | 6.1 |
| DRAXIN | 16.7 | 0.8 | 4.2 | 13.1 |
| ANGPT2 | 16.0 | 0.8 | 4.8 | 13.3 |
| PTX3 | 14.9 | 0.7 | 2.3 | 10.7 |
| PLAU | 14.1 | 0.9 | 3.1 | 12.7 |
| WNT5A | 13.4 | 1.2 | 7.3 | 16.0 |
| IGFBP5 | 12.4 | 21.7 | 85.9 | 267.7 |
| CCL3 | 11.0 | 0.9 | 2.2 | 10.0 |
| NTN1 | 9.0 | 2.8 | 11.2 | 25.5 |
| VEGFA | 8.8 | 19.4 | 34.0 | 169.9 |
| SEMA3A | 7.5 | 1.0 | 2.3 | 7.3 |
| CYTL1 | 6.8 | 0.9 | 2.5 | 6.2 |
| RGMA | 6.4 | 16.7 | 45.4 | 106.8 |
| FBLN1 | 6.3 | 6.6 | 11.7 | 41.5 |
| HGF | 5.6 | 1.2 | 2.3 | 6.7 |
| HMGB1 | 5.4 | 40.2 | 99.3 | 217.9 |
| PLAT | 5.0 | 9.5 | 13.9 | 47.8 |
| SEMA3C | 4.9 | 1.9 | 3.5 | 9.4 |
| PDGFC | 4.8 | 2.9 | 6.4 | 14.1 |
| PGF | 4.5 | 3.0 | 4.5 | 13.6 |
| IGFBP7 | 4.3 | 77.0 | 113.2 | 331.6 |
| TGFB2 | 3.9 | 6.1 | 12.9 | 23.9 |
| GRN | 3.9 | 23.4 | 46.3 | 90.7 |
| CXCL12 | 3.7 | 3.4 | 4.7 | 12.6 |
| TGFB1 | 3.1 | 9.2 | 14.7 | 28.2 |

Grey: CXCR4 ligand
Blue: MCIP chemoattractants, includes CXCL12

Interestingly, glioblastoma also secrete other factors known to chemoattract MCIPs; namely CXCL12, HGF, and NRG1 all scored in our analysis (Fig 1A and Table 1). Given that HGGs highly express several chemotactic factors crucial for MCIP migration, we postulated that MCIPs would exhibit targeted migration to HGGs.

**Figure 1:**
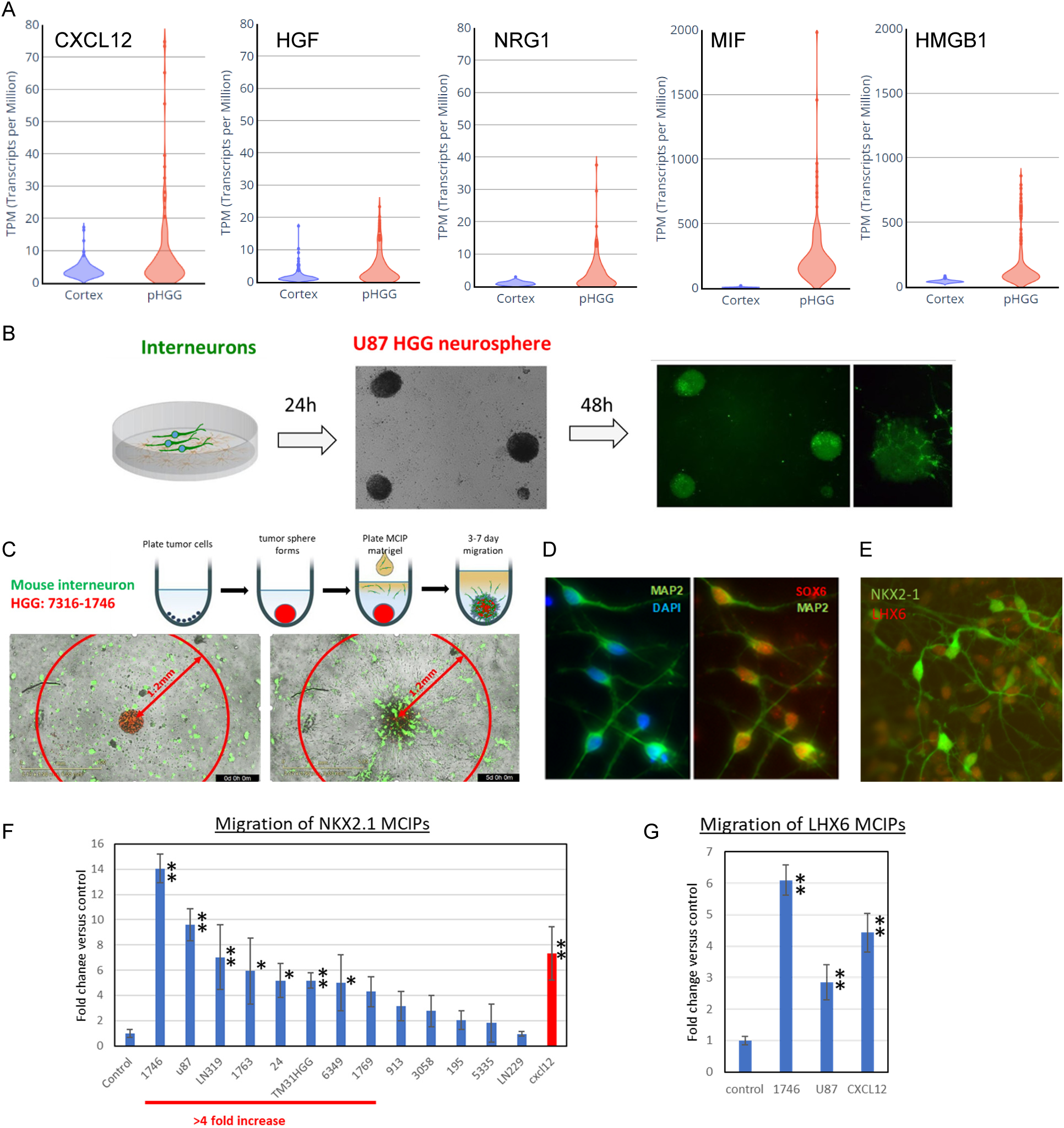
High-grade glioma chemoattract MCIPs in vitro. (A) Violin plots comparing expression (transcripts per million, TPM) of known MCIP chemoattractants and CXCR4 ligands in pHGG and normal cortex. (B) Mouse MGE derived interneurons (green) are attracted to U87 glioblastoma spheres: within 48h the U87 glioma spheres become GFP positive denoting MCIP honing. (C) Mouse MGE derived interneurons (green) are attracted to a 7316-1746 HGG sphere (red) in a spheroid migration assay when the glioma spheres are encased in a Matrigel matrix containing MCIPs (green). MCIPs are allowed to migrate for 5 days. A tumor sphere that induces an MCIP migration front with a radius of >0.5mm is considered as attracting MCIPs. Images at day 0 and 5 post plating. (D) Human MCIP IF Staining with MCIP Markers MAP2 (green) and SOX6 (red). (E) Human MCIP IF Staining with MCIP Markers LHX6 and NKX2-1. As expected at the 4 week timepoint shown, cells are NKX2-1 positive (red), while a subset is positive already for LHX6 (green). (F-G). Incucyte transwell migration: NKX2.1 MCIPs (F) or LHX6 MCIPs (G) are plated in the top well. 24h later a chemoattractant (chemokine or conditioned media) is added to the bottom well and migration is monitored. Data are plotted as fold change number of cells on the bottom of the membrane compared to control after 3 days. Unconditioned media serves as a control. *p<0.02, **p<0.005

### MCIPs potently migrate to glioma cells *in vitro*

Given the fact that several canonical MCIP chemoattractants are secreted by HGG, we evaluated the ability of both mouse and human MCIPs to migrate to HGG. To test this hypothesis, murine MGE cells were harvested from E13.5 embryos that express Green Fluorescent Protein (GFP), dissociated, and plated on Matrigel coated culture dishes. These dissociated cells quickly differentiate to MCIPs in culture. The following day, human HGG spheroids, expressing nuclear red fluorescent protein (RFP) were plated on top of the MGE derived interneurons and live imaging was performed every 6 hours using IncuCyte^TM^. Robust MCIP migration to the glioma spheres was observed within 48 hours of co-culture (U87: Figure 1B, 7316-1746: Video 1). To standardize our approach, we also evaluated the ability of MCIPs to migrate to glioblastoma in a spheroid migration assay, where glioblastoma spheres (RFP) are encased in Matrigel containing MCIPs (GFP). Any HGG sphere that induced an MCIP migration front with a diameter of >0.5mm in 5 days (Figure 1C and Video 2) was considered a success. Of the 5 lines evaluated, 4 met our criteria and exhibited mouse MCIP chemoattraction. Migration was observed in lines 7316-1746, U87, 7316-1763 and 7316-3058, but not in line 7316-1769.

To ascertain if these results could be recapitulated with human MCIPs, embryonic stem cells (ES cells) were chemically patterned to generate MGE progenitor cells followed by GABAergic cortical interneurons over 4-5 weeks. Cells were rendered post-mitotic using a combination of DAPT and CDK4/6 inhibitor. Cellular identity and appropriate progression through differentiation was verified on a weekly basis via immunocytochemistry for MAP2, SOX6, NKX2-1 and LHX6 (Figure 1D, E). As expected, we see 100% positivity for MAP2 and SOX6. NKX2-1 expression levels are initially high, but NKX2.1 becomes downregulated as MCIPs mature, exit the cell cycle, and upregulate LHX6. This cell cycle exit, with LHX6 upregulation, happens stochastically during the differentiation process, with a gradual increase in the percentage of LHX6 positive cells. The staining in Figure 1E shows a mixture of NKX2.1 positive cells and LHX6 positive cells. We routinely derive MCIPs from ES line HES3, which expresses GFP under the NKX2.1 promotor (hereby called NKX2.1 line), and from ES line H9 modified to express Citrine (GFP) under the LHX6 promotor (hereby called the LHX6 line).

We then compared the ability of different glioblastoma lines to chemoattract MCIPs using a transwell migration assay, where 20,000 MCIPs per well were plated in the upper chamber of Matrigel-coated 96 well transwell migration plate and allowed to adhere overnight. The following day, HGG conditioned MCIP media was added in the lower chamber of the transwell plate. Fresh MCIP base media was used as a negative control and CXCL12, which has been shown to induce interneuron migration from the MGE, was used as a positive control. We calculated the fold change in migration by dividing the number of MCIPs that reached the bottom well in conditioned versus control media. A four-fold increase in NKX2.1 MCIP migration to the bottom well was observed for 8 of the 13 conditioned HGG medias evaluated (ANOVA p-value = 2x10^-10^) as shown in Figure 1F. For 7 of these HGG lines this increase in migration was significant (Tukey Kramer p-value<0.02), for the 8^th^ HGG line significance was borderline (p=0.07, Table 2). We sought to determine if these results were generalizable across MCIPs derived from other cell lines as this could allow for either patient-derived MCIP transplant or for donor cells to be HLA-matched and used off the shelf to avoid immunosuppression with transplantation in the clinic. We observed that LHX6 MCIPs were similarly able to migrate to high-grade glioma conditioned media (Figure 1G, tumor conditioned media from 7316-1746 and U87, t-test p-value <0.005).

**Table 2:** Fold change migration of NKX2.1 MCIPs towards tumor conditioned media.

| Conditioned media | Average fold vs control | SD | p-value Tukey-Kramer test |
| --- | --- | --- | --- |
| Control | 1.00 | 0.31 | - |
| 7316-1746 | 14.06 | 1.14 | 2.2E-10 |
| U87 | 9.62 | 1.27 | 8.3E-08 |
| LN319 | 7.03 | 2.56 | 6.0E-05 |
| 7316-1763 | 5.94 | 2.63 | 6.9E-03 |
| TM31HGG | 5.50 | 0.32 | 1.8E-02 |
| 7316-24 | 5.18 | 1.36 | 8.5E-03 |
| 7316-6349 | 5.00 | 2.20 | 1.4E-02 |
| 7316-1769 | 4.31 | 1.19 | 7.2E-02 |
| 7316-913 | 3.15 | 1.18 | 5.7E-01 |
| 7316-3058 | 2.77 | 1.25 | 8.1E-01 |
| 7316-5335 | 2.50 | 1.27 | 9.7E-01 |
| 7316-195 | 2.05 | 0.76 | 9.9E-01 |
| LN229 | 0.95 | 0.19 | 1.0E+00 |
| cxcl12 (100ng/ml) | 8.37 | 1.46 | 2.5E-05 |

### MCIPs potently migrate to HGG cells *in vivo*

#### Mouse MCIP migration

In order to test the migratory capacity of MCIPs to HGG *in vivo*, U87 cells were orthotopically xenografted in the frontal lobe of nude mice and allowed to establish tumors over the course of a week. Murine MCIPs harvested from the MGE region of RFP-expressing mouse embryos were grafted approximately 2 mm apart from HGG cells and allowed to migrate for seven days post-transplant. As demonstrated in Figure 2A, murine MCIPs potently migrated to the glioma, not only reaching the edge of the main neoplasm (Figure 2B), but also encapsulating satellite lesions (Figure 2C) suggesting MCIPs possess powerful honing capabilities for infiltrative tumor.

**Figure 2:**
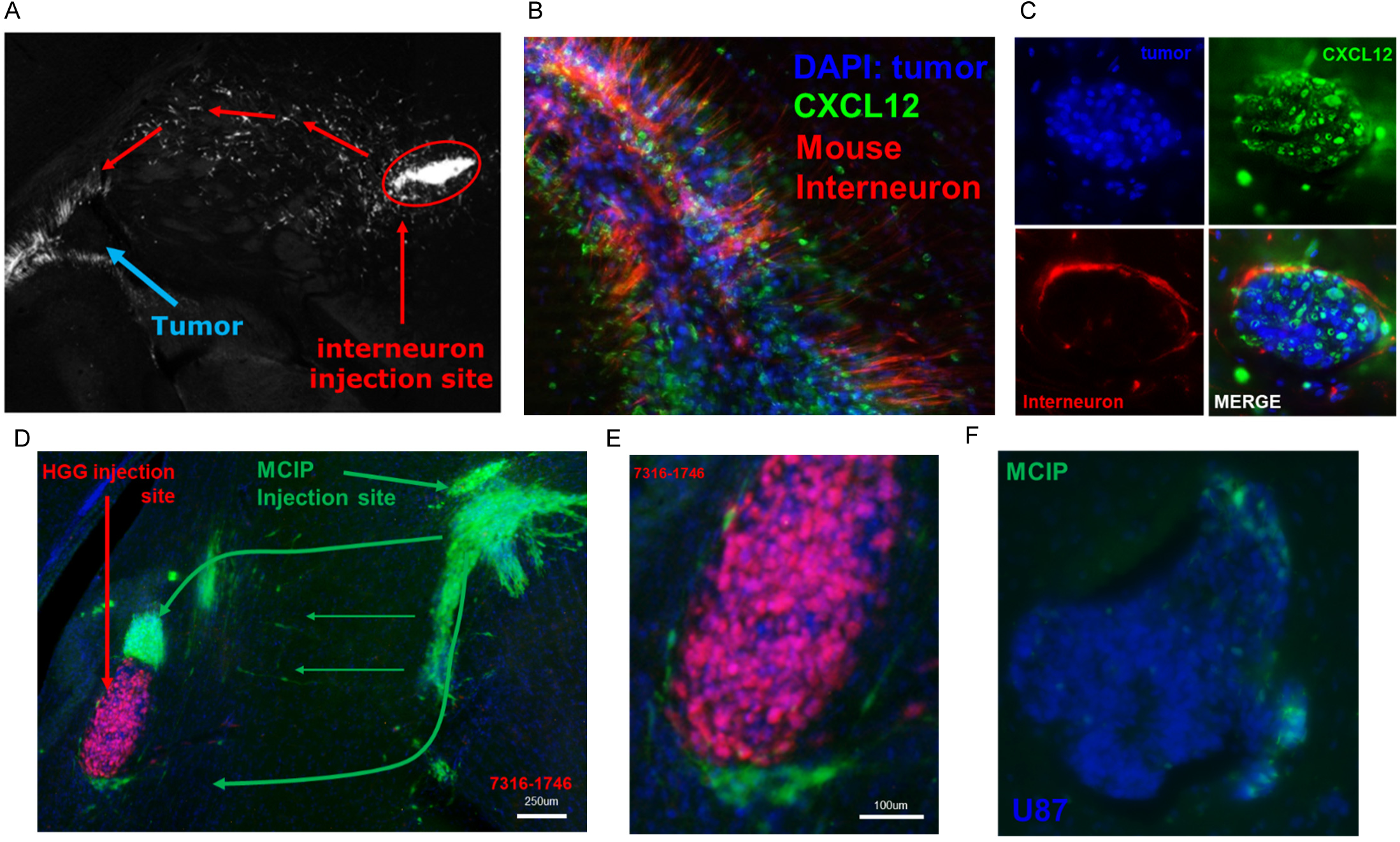
High-grade glioma chemoattract MCIPs in vivo: (A-C) Mouse MCIPs migrate to the site of a glioblastoma (U87). (A) Low magnification image (4x) showing an interneuron stream from the MCIP injection site (200k cells) to the tumor site (100k cells). (B) MCIPs (red) arrive at the location of the tumor (blue, U87) in 7 days and wrap around the main glioblastoma (green=CXCL12). (C) Micro-lesion of U87 wrapped in MCIPs (red) (green=CXCL12). (D-F) NKX2.1 Human MCIPs (green) migrate to the site of a glioblastoma within 14 days when injected 2 mm away: (D) Low (4x) magnification of HGG 7316-1746 (red) and MCIP injection (green, arrow) sites and stream of MCIPs reaching the tumor. (E) Magnification showing MCIPs wrapping around the 7316-1746 tumor (red), similar to the mouse MCIP experiments. (F) Human MCIPs (green) that have reached the margin of U87 (Blue).

#### Human MCIP migration

Mice were orthotopically injected with HGG lines 7316-1746, U87 or LN229. After 7 days, human NKX2.1 MCIPs, which express the Citrine fluorescent marker, were injected at 2 mm distance. Animals were sacrificed 14 days post implantation and migration was visualized by staining for Citrine (GFP). Implanted MCIPs effectively targeted 7316-1746 cells as well as U87s *in vivo* over a two-week span, but not LN229 HGG cells (Figure 2D-F).

### EGFR-BiTE expressing MCIPs promote potent glioma death *in vitro*

We next examined if MCIPs could be neuro-engineered to locally secrete a therapeutic agent. As proof of principle, we stably transduced LHX6 MCIPs to express a Bispecific T-cell Engager (BiTE) targeting wild-type EGFR or a non-targeting CD19 control. We used CD19 as a control because it is minimally expressed in brain. BiTEs are essentially two antibody fragments (scFv or single-chain fragment variable) connected by a linker which allow a T-cell to be connected to the tumor cell via its T-cell receptor (CD3). When both sides of the BiTE are engaged, the T-cell becomes activated, releasing perforin and granzyme and clearing the glioma cell. This allows T-cells to recognize targeted tumor cells that had previously evaded the immune system.^18,19^ Figure 3A gives an overview of the BiTE composition. MCIP BiTE expression was confirmed using qPCR. To evaluate the efficacy of our BiTE we initially transfected our constructs into 293Ts and co-plated them with tumor cells and T-cells and monitored tumor cell killing using the Incucyte live imager. The cytotox green reagent was added to visualize dead cells (GFP positive). Video 3 and 4 show potent cell killing induced by the targeting EGFR-BiTE but not the control CD19-BiTE. Figure 3B shows a graph of the induced killing.

**Figure 3:**
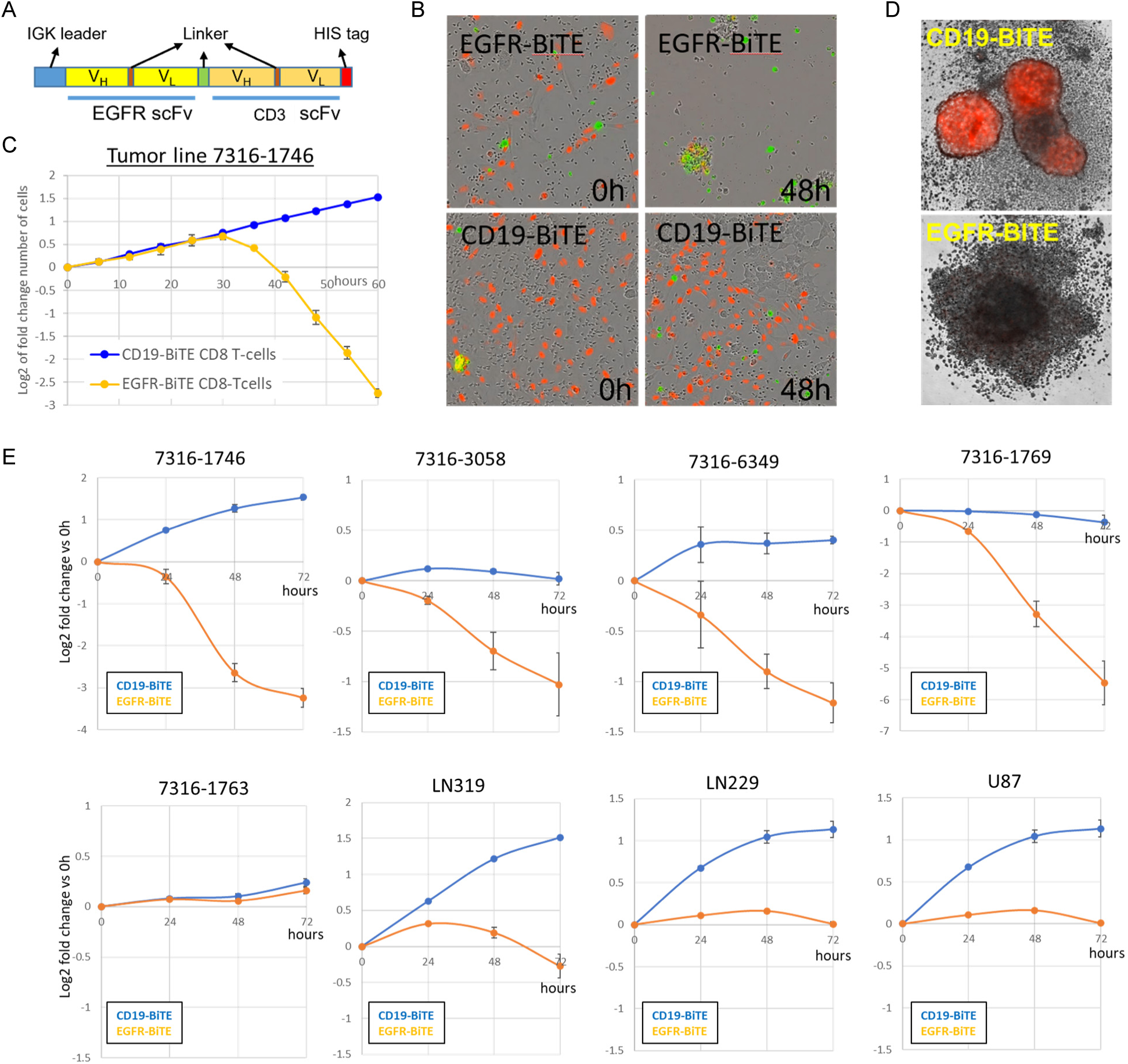
BiTE expressing MCIPs potently kill glioblastoma *in vitro*: (A) Schematic overview of the BiTE composition, with IGK leader sequence to stimulate secretion, anti-EGFR and anti-CD3 scFvs and linker sequences. (B-C). An EGFR BiTE potently eliminates glioblastoma line 7316-1746 *in vitro*: (B) 2D killing assay (0 and 48h) showing 1746 pHGG cells (red, 5k) mixed with targeting (EGFR) or control (CD19) BiTE expressing 293T cells (2k) and CD8 T-cells (50k). A Cytotox agent stains dead cells green. (C) Graphical representation of 3B plotted as the log2 of the fold change compared to the number of cells present at hour 0. (D) Images of 3058 pHGG sphere (red, 5k) mixed with BiTE expressing MCIPs (2k) and CD8 T-cells (50k, small cells surrounding sphere), show potent tumor killing with EGFR-BiTE at 48h. A CD19 control BiTE did not kill. (E) BiTE killing assay: Graphical representation of the survival of glioblastoma cell lines mixed with targeting (EGFR) or control (CD19) BiTE expressing MCIPs and CD8 T-cells. The log2 of the fold change compared to the number of cells present at hour 0 is plotted. Significant cell death is induced in 4/8 lines evaluated. All lines except 7316-1763 exhibit a statistically significant change (p<0.005, t-test)

To determine if BiTE-expressing MCIPs were capable of clearing HGG cells, MCIPs expressing either an EGFR BiTE or CD19 control BiTE were co-plated with HGG cells at a ratio of 0.1:1. Figure 3D shows an image of the killing induced by CD8 T-cells when the MCIPs expressing EGFR-BiTE but not the CD19-BiTE are co-cultured in the 7316-3058 spheroid line. Potent HGG killing (>50% cell death in 72h) was observed in 4/8 of the cell lines tested and reduced growth was observed in 3/8 lines tested irrespective of the effector to target ratio (Figure 3E). Minimal to no glioma cell death was observed in 1/8 cell lines tested.

### MCIPs expressing an EGFR-BiTE delay glioma onset and increase survival in pediatric high-grade glioma xenograft models

Next, we evaluated the efficacy of the MCIP-based therapy *in vivo* using pediatric HGG xenograft models. Since sequential transplant of HGG cells, MCIPs, and intraventricular cannulation to deliver exogenous T-cells to immunocompromised mice would require a series of three surgeries within a three-week window, we decided to perform pilot experiments with co-transplantation of the aforementioned cells in a single orthotopic injection in 9 mice per treatment arm. Tumor size was monitored weekly via live chemiluminescence imaging. Mice that received EGFR-BiTE expressing MCIPs demonstrated delayed tumor onset and maintained a subthreshold luciferase signal at 8 weeks post-transplant of 7316-3058 thalamic midline glioma cells (Figure 4A). In contrast, mice treated with the CD19 BiTE all had robust luciferase signal demonstrating significant tumor growth. Furthermore, this corresponded with significantly improved median overall survival for EGFR-BiTE treated mice (Figure 4B), with subset of mice doubling their overall survival (p=0.02). Unfortunately, all mice in the EGFR BiTE cohort did ultimately develop tumors, which were detectable on live imaging and proved lethal. We then repeated this with the 7316-6349 midline glioma line which was orthotopically grafted within the pons. Again, we observed that mice treated with EGFR-BiTE expressing MCIPs had prolonged survival (p=0.025) when compared to CD19 BiTE expressing controls (Figure 4C).

**Figure 4:**
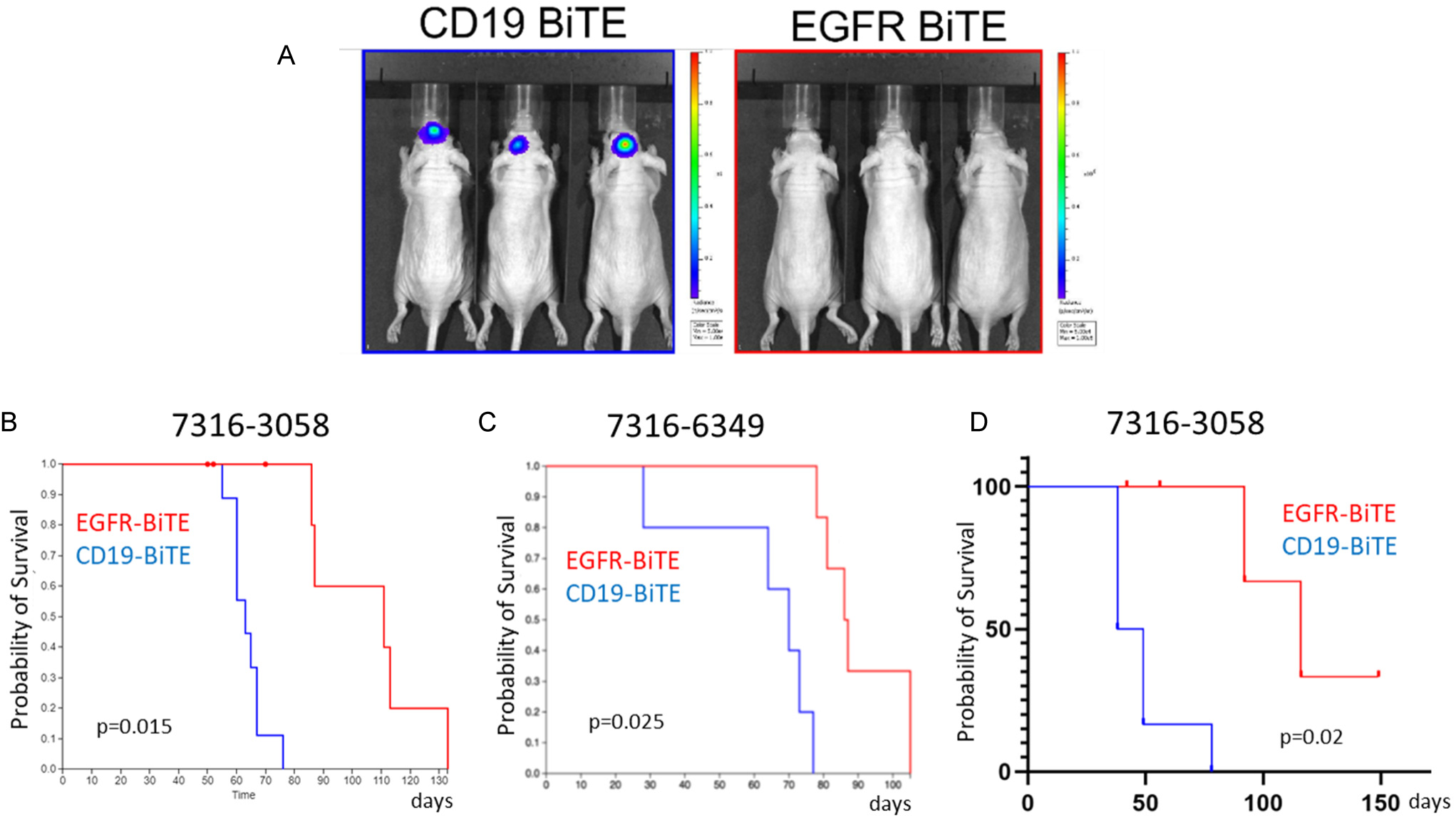
BiTE expressing MCIPs enhance survival of mice with high-grade glioma: (A-B) Mice were injected with 7316-3058 tumor cells (500k cells), BiTE expressing MCIPs (100k cells) and T-cells (10^6^ cells) in a single location. CD19-BiTE was used as a control. (A) Luminesce signal shows a tumor present with CD19-BiTE MCIP controls, the signal is below detection threshold with EGFR-BiTE MCIPs. (14d post injection) (B) Survival curve of the 7316-3058 xenografts showing that the EGFR-BiTE treated mice have an extended survival (co-injection tumor cells/MCIPs/T-cells 9 mice per study arm, p=0.015). (C) Survival curve of the 7316-6349 xenografts showing that the EGFR-BiTE treated mice have an extended survival (co-injection tumor cells/MCIPs/T-cells 6 mice per study arm, p=0.025). Mice were injected with 7316-6349 tumor cells (500k cells), BiTE expressing MCIPs (100k cells) and T-cells (10^6^ cells) in a single location. CD19-BiTE was used as a control. (D) Survival curve of the 7316-3058 xenografts showing that the EGFR-BiTE treated mice have an extended survival (co-injection tumor cells/MCIPs 6 mice per study arm, p=0.02) when we first allowed the tumor to establish before placing a cannula in the ventricles and supplying CD8 T-cells. CD19-BiTE was used as a control.

Given that CD8 T-cells are only capable of expansion and activation for approximately one month prior to exhaustion within the brain, we questioned if recurrent T-cell dosing might amplify the anti-tumor response. To test this, we orthotopically allografted 7316-3058 midline glioma cells in the thalamus of nude mice along with EGFR or CD19 BiTE expressing MCIPs with 6 mice per treatment arm. Tumors were allowed to establish over a three-week period to ensure consistent signals above threshold were observed on live imaging. Mice then underwent implantation of an intraventricular cannula to allow for weekly administration of 2.5 million T-cells over the course of four weeks. Tumor size was monitored via luciferase signal on IVIS live imaging. Two mice were censored due to complications from T-cell infusion. Consistent with the co-injections, mice with established brain tumors treated with EGFR-BiTE expressing MCIPs showed reduction in luciferase signal with repeated T-cell dosing and had extended survival (p=0.02) compared to CD19 BiTE controls (Figure 4D). Furthermore, one of the mice in the EGFR-BiTE cohort fully cleared its tumor with no evidence of residual tumor on necropsy five months after initial xenografting. Collectively, these proof of principle experiments suggest that MCIP-delivered therapies can kill HGG cells *in vivo*.

## Discussion

Despite significant advances in our knowledge of the cellular and molecular drivers of HGG, these tumors remain almost universally fatal^1^. For glioblastoma, the standard of care, which consists of maximal safe surgical resection of the tumor followed by adjuvant chemotherapy, has not evolved since 2005^20,21^. Although some targeted, cellular, and immunotherapies have shown promise in the clinic, most fail to make a substantial impact on the survival of patients. The inability of many agents to reach the glioma in high enough concentrations or numbers remains a bottleneck.^4^ Cellular vectors, like the presented MCIPs can materially change our ability to deliver those therapeutic agents to an HGG.

Given that MCIPs display tropism for HGGs and are capable of delivering a therapeutic cargo, we envision that administration of MCIP-based therapies could be incorporated into standard-of-care surgery, or at time of recurrence, with injection of MCIPs into the resection cavity in order to clear residual, highly infiltrative cells. However, MCIPs could also be directly injected into or near residual tumor for patients in which a gross-total resection is not possible, though the minimum number of MCIPs required to effectively clear a given tumor also remains unknown. Molecularly engineered MCIPs can be equipped in a multitude of ways including secretion of an enzyme for prodrugs which otherwise might not cross the blood brain barrier, delivery of BiTEs, secretion of anti-tumor antibodies, secretion of peptides modulating the tumor microenvironment, and delivery of oncolytic viruses among others.

Several BiTE therapies are approved by the FDA, including Blinatumomab for patients non-Hodgkin Lymphoma^22^. While direct injection of BiTEs into the brain could be considered for HGG, there are several limitations to this approach. BiTEs have a very short half-life (2h)^23^;Blinatumomab for example requires continuous infusion for 28 days to achieve adequate dosing to target tissues^24^. For HGG, this is extraordinarily challenging as continuous access to the brain via an Ommaya reservoir poses a high risk of infection. Furthermore, BiTEs also have limited diffusion capabilities and may not reach the tumor through intraventricular delivery. In contrast, MCIPs continuously secrete BiTEs locally to the tumor, which also can limit on-target, off-tumor effects and might broaden the antigenic targets by allowing the use of antigens that are unique to brain but not to other tissues. While we describe the use EGFR-BiTEs which are non-unique antigen for our proof of principle experiments, we anticipate that initial trials involving BiTE delivery would utilize a unique target to minimize risk of additional toxicities.

Notably, there is precedent for a parallel approach using fetal neural stem cells as a delivery vector in glioblastoma for which several clinical trials are ongoing (NCT01478852, NCT03072134, NCT05139056)^25–29^. In contrast to neural stem cells which have been shown to undergo a period of expansion for the first 14 days post-injection, MCIPs are post-mitotic and incapable of expansion post-transplant, mitigating concerns for aberrant proliferation and potential tumor formation. MCIPs also are capable of long-term survival following transplant with persistence of at least 5 months after initial transplant. This is substantially longer than that observed with neural stem cells, the majority of which persist less than 30 days post-injection and with only a few cells observed 79 days post-transplant in one post-mortem sample^29^. As such, MCIPs provide a more stable delivery system that may require less frequent dosing than neural stem cells to achieve adequate treatment response. The need for frequent recurrent dosing has been a substantial limitation of neural stem cells in current trials as repeated intratumoral dosing via catheter has been severely limited by scarring at the catheter site^27^. Furthermore, MCIPs can be derived from iPSCs^15^, circumventing both the ethical concerns regarding the use of fetal cells in therapies as well as the potential need for immunosuppression with transplant, as cells could be HLA-matched to be administered off the shelf or generated from individual patient samples.

MCIPs-based therapies also diverge from other cellular therapies as these cells are native to the brain and can functionally integrate. Recent studies have shown that glutamatergic signaling via AMPA-receptor mediated synapses between neurons and glioma cells drives HGG proliferation and progression^30^. Furthermore, glutamate-driven excitotoxicity has been implicated in the peritumoral loss of fast-spiking parvalbumin positive interneurons in orthotopic xenograft models, which subsequently produces cortical network disinhibition and peritumoral hyperexcitability, allowing for further tumor progression^30^. This raises the question if MCIP-based therapies could also have the secondary benefit of modulating aberrant neuronal-glial communication contributing to the underlying disease pathophysiology and significant patient morbidity. Enticingly, there is precedent for human MCIP transplantation, with an ongoing clinical trial evaluating the safety and efficacy of interneuron transplant for the treatment of medically refractory epilepsy (NCT05135091). Further evaluation of this potential in xenograft models is required.

There are potential limitations to MCIP-based therapies such as the presence of multifocal, bulky disease which may reduce the efficacy of treatment by diluting the dosage of MCIPs reaching each lesion. The optimal method of MCIP delivery remains a key question as intraventricular rather than intracerebral injection would allow for repeated dosing of MCIPs to enhance response given that MCIPs are only capable of migration for approximately 1 month after transplant^9^. Additional studies are also required to determine if MCIPs retain their migratory capacity and survive with standard of care therapy. Finally, we have shown that the majority of glioblastoma and HGG attract these MCIPs *in vitro*; however, the molecular mechanisms mediating MCIP-to-HGG migration have yet to be elucidated which is pertinent to identifying the targeted population eligible for MCIP-based therapies.

In conclusion, we have demonstrated that MCIPs have significant honing capabilities for HGG both *in vitro* and *in vivo*. Crucially, this honing is independent of antigens or the tumor microenvironment and MCIPs generated from different sources all have the capacity to migrate to these tumors. We showed that MCIPs can be neuro-engineered to deliver anti-tumor cargos, with our example of an EGFR-BiTE being just a proof of principle. As injected MCIPs will not leave the brain and locally deliver a therapy, therapeutic proteins that would otherwise induce systemic toxicity can be considered as a cargo.

## Materials and methods

### *In silico* analysis of pHGG secretome

We used several datasets to analyze the pHGG secretome involved in migration. We used normalized (TPM) RNAseq data from 133 pHGG tumors (CBTN, pedcbio.org) and normal cortex (GTEX database, gtexportal.org). First, we selected all genes that were known to be secreted (1891 genes, The Human Protein Atlas secreted proteins, https://www.proteinatlas.org/search/protein_class%3APredicted+secreted+proteins). Next, we determined the median TPM value in normal cortex and the 80^th^ percentile expression across the 133 pHGG samples (which corresponds to the expression level of a specific gene in the tumor that has the 20% highest expression) and determined the fold change between the 80^th^ percentile pHGG expression and median expression in cortex. We used the 80^th^ percentile expression to cast a wide net and because we considered chemoattraction to 1/5 tumors worthwhile pursuing. We only considered a gene of interest when the determined fold change was at least 3. Finally, we manually curated the resulting gene list for known genes involved in chemoattraction and resulted in the 38 genes listed in Table 1.

### Reagents

Antibodies: LHX6 (Sc-271433, SCBT), SOX6 (AB5805, Millipore), anti-GFP (Ab13970, Abcam), NKX2.1 (Ab76013, Abcam), MAP2 (Ab5392, Abcam), CXCL12 (MAB350, R&D systems). BiTEs: EGFR and CD19 bi-specific T-cell engagers were generated as previously described^31^ by the gene synthesis service of Genscript and cloned in a lentiviral construct under the CAG promotor.

The CXCL12 chemokine was purchased from R&D Systems (350-NS-050).

### Glioma cell culture

Human pediatric high-grade glioma stem cell lines (7316-24, 7316-195, 7316-913, 7316-1746, 7316-1763, 7316-1769, 7316-3058, 7316-5335, and 7316-6949) were acquired from the Children’s Brain Tumor Network and were characterized as previously described^32^ and maintained in glioma stem cell media (DMEM/F12 medium supplemented with 1% GlutaMAX (Gibco, Billings, MT), 20 ng/mL basic fibroblast growth factor (PeproTech, Rocky Hill, NJ), 20 ng/mL epidermal growth factor (StemCell Technologies, Vancouver, Canada), 1x B-27 supplement (Thermo-Fisher Scientific, Waltham, MA), 1X N-2 supplement (Thermo-Fisher Scientific, Waltham, MA), 4 µg/mL heparin (StemCell Technologies, Vancouver, Canada), and 100 U/mL penicillin-streptomycin (Life Technologies, Carlsbad, CA). Adult glioblastoma lines were maintained in DMEM supplemented with 10% FBS (Gemini Bio-Products, West Sacramento, CA), and 100U/mL penicillin-streptomycin (LN229, LN319, U87). The human adult glioblastoma TM31HGG was dedifferentiated from the serum grown TM31 cell line (Received from the RiKEN biorepository) by propagating it in the glioma stem cell media described above. All glioma lines were stably transduced with a lentivirus expressing nuclear red-fluorescent protein (NucLight Red, Sartorius) or with lentivirus expressing green-fluorescent protein and luciferase for imaging purposes.

### MCIP culture and differentiation

MCIPs were differentiated from human embryonic stem cells (NKX2.1-GFP and LHX6 Citrine)^33,34^ using a modified version of the protocol from Ni *et al.* 2019 and 2020^35,36^ The stem cells were cultured on Matrigel (Corning 1:50) coated plated in StemMACS™ iPS-Brew XF media (Miltenyi Biotec) supplemented with 10 µM Y27632 ROCK inhibitor (ApexBio).

Stem cells were dissociated with Accutase (Sigma-Aldrich) and replated on low attachment plates (Corning) into SRM media consisting of DMEM (Invitrogen),15% knockout serum replacement (Invitrogen), 2 mM L-glutamine (Gibco), β-mercaptoethanol (Gibco) and 10 µM Y27632 ROCK inhibitor (ApexBio).

From day 2-7 cells were cultured in SRM media supplemented with 0.1 µM LDN193189 (Tocris Biosciences), 10 µM SB431452(Tocris Biosciences), 0.1 µM SAG (Tocris Biosciences), and 5 µM IWP2 (Selleckchem). Media was changed every other day. From day 8-14 cells were cultured in SRM media supplemented with 0.1 µM LDN193189 (Tocris Biosciences) and 0.1 µM SAG (Tocris Biosciences). From day 15-21 the cells were cultured in N2AA media consisting of DMEM/F12 media supplemented with 0.5% N2 supplement (Sigma-Aldrich), 200 µM AA-ascorbic acid (Sigma-Aldrich), 0.1 µM SAG (Tocris Biosciences), and 50 ng/mL FGF8 (Prospec Bio). After day 21 cells were cultured in N2AA media supplemented with 2 U/mL Turbo DNase (Thermo Fisher), 10 µg/mL GDNF (Thermo Fisher), 50 µg/mL BDNF (Thermo Fisher), B27 (Thermo Fisher) and 2uM MEK inhibitor (PD-0325901 Tocris Biosciences). One week prior to dissociation cells were treated with mitotic inhibitors 10 mM DAPT, 5 mM PD 0332991 and MEK inhibitor. The MCIP organoids were dissociated using 0.05% trypsin (Thermo Fisher) and 100 mM trehalose (Thermo Fisher) and were fully triturated. Trypsin was neutralized with N2AA media supplemented with 2 U/mL Turbo DNase (Thermo Fisher) 10 µg/mL GDNF (Thermo Fisher), 50 µg/mL BDNF (Thermo Fisher), B27 (Thermo Fisher), and MEK inhibitor (PD0325901 2uM; 4192 Tocris). Dissociated cells were plated on Matrigel (Corning) coated plates for 24 to 48 hours. Finally, cells were chemical dissociation to single cell mixtures with TrypLE (Thermo Fisher) and exposed to 5ml N2AA media 1:10,000 Benzoase (Sigma-Aldrich) for 5min and replated using N2AA media.

For therapeutic assays, WA-09 cells were stably transduced with lentivirus constructs expressing an EGFR BiTE or CD19 control BiTE.

### T-cell expansion

Human CD8 T-cells were obtained from the University of Pennsylvania Human Immunology Core and expanded as previously described^37^.

### IncuCyte™ Spheroid Migration Assay

Glioma cell lines were plated in triplicate at 5000 cells per well in 100 µL of N2AA media in 96-well ultra-low attachment plates (S-bio). Plates were then spun at 125g for 5 minutes to promote spheroid formation. The following day, 50,000 cortical interneurons in 45 µL of N2AA media were mixed with 45 µL of growth factor reduced Matrigel (Corning) (4C). After cooling the glioma sphere plate to 4C for 5min, the 90 µL MCIP mixture was gently added on top of each glioma sphere containing well and allowed to solidify for 30min at 37C. Plates were scanned in the Incucyte with the spheroid model every 8 hours for 5 days.

### IncuCyte™ Transwell Migration Assay

Cells were plated in MCIP base media supplemented with BDNF and GDNF. Transwell migration plates (Sartorius) were coated with growth factor depleted Matrigel (Corning) diluted 1:50 in DMEM. After a minimum of 3-hour incubation at 37C, dissociated MCIPs were plated at 20,000 cells per filter in 60ul media and 200ul media was plated in the bottom well. The following day, 15ul of the top well was replaced with fresh BDNF/GDNF supplemented media. Next, 150uL the media in the lower chamber of the transwell migration plate was removed, 50ul fresh BDNF/GDNF supplemented media was added as well as 100ul MCIP base media, glioma-conditioned base media, or CXCL12 conditioned media (100ng/ml final concentration). Plates were imaged using IncuCyte™ live imaging software every 4 hours for a total of 5 days. Glioma-conditioned media was generated by incubating two million HGG cells in 10mL of MCIP base media for 48 hours.

### 2-Dimensional IncuCyte™ Killing Assays

HGG cell lines were plated at 10,000 cells per well and MCIPs were plated at 2,000 cells per well in 96 well plates and allowed to adhere overnight. The following day, 50,000 human CD8 T-cells were introduced to each well. Plates were then monitored for 3 days using the IncuCyte software.

### 3-Dimensional IncuCyte™ Killing Assays

A total of 5,000 HGG cells and 1,000 MCIPs were plated in ultra-low attachment round bottom 96-well plates (S-bio) and spun down at 125g for 5 minutes to promote spheroid formation. The following day, 10,000 human CD8 T-cells were introduced to each well. Plates were then scanned for 3 days using the IncuCyte software.

### Animals

All *in vivo* experiments were approved by the Children’s Hospital of Philadelphia Institutional Care and Use Committee. Animals were housed per institution guidelines in a vivarium with *ad libitum* food and water access and a 12-hour light/dark cycle.

### Medial Ganglionic Eminence Harvest

Donor MGE cells were generated by breeding homozygous GFP/RFP mice to wild type females on a C57BL/6J background. Embryonic day 0.5 (E0.5) was defined by the time at which vaginal sperm plug was detected. Embryonic MGE cells were dissected at E13.5 in Neurobasal media (Gibco) supplemented by 1% B27 (Thermo-Fisher Scientific, Waltham, MA). For dissociation, DNAse was added at 1U/uL and cells were mechanically dissociated to a single-cell suspension using a flamed Pasteur pipette, then collected by centrifugation (500 rpm for 5 minutes at 4 °C).

### Murine Spheroid Migration Assay

Murine MGE cells were plated on Matrigel coated plates at a density of 100,000 cells/cm^2^ in Neurobasal media supplemented with 1% B27 and allowed to adhere overnight. The following day, human HGG spheroids were plated in each well and then imaged for 72 hours on the IncuCyte™ to assess for migration.

### Orthotopic MGE Allografts and Glioma Xenografts

At postnatal day 1, Nu/J mice (Jackson labs) were anesthetized by reducing their body temperature on ice and injected with 100,000 glioma cells. Pups are aged and a total of 50,000 donor murine MGE cells were injected in 20-day old pups using a Hamilton syringe. MGE cells were allowed to migrate over the course of 7 days prior to brain harvest for histology.

### Glioma and Interneuron Xenografts

Eight to 10-week-old Nu/J mice were anesthetized with 2.5% isoflurane, placed into the stereotactic frame, then injected sequentially with 500,000 glioma cells. Tumors were allowed to establish for 1 week, then mice underwent a second surgical procedure with stereotactic injection of 100,000 MCIPs a total of 2 mm apart from initial injection site. Stereotactic coordinates used to target the cortex for tumor or interneurons, respectively were: 1 mm anterior to bregma, 2 mm lateral to bregma and 1mm depth from pia (glioma) or -0.4 mm posterior to bregma, 3 mm lateral to bregma, and 1 mm depth from pia (MCIPs). For striatal interneuron injections, stereotactic coordinates used were 1mm anterior to bregma, 2mm lateral to bregma, and 3.5 mm depth from pia (glioma) or 0.7mm posterior to bregma, 3mm lateral to bregma, and 3.5 mm depth from pia. MCIPs were allowed to migrate for two weeks prior to harvesting brains for histology.

For *in vivo* killing assays, an admixture of 200,000 either CD19 control BiTE or EGFR-BiTE expressing MCIPs and 500,000 glioma cells were co-injected at 1 mm posterior to bregma, 0.8 mm lateral to bregma, and 3.5 mm depth from skull (7816-3058, thalamus) or 0.8 mm posterior to bregma, 1.00 mm lateral to bregma, and 5 mm depth from skull (7816-6349, pons). CD8 T-cells were obtained and expanded as above for all *in vivo* experiments. All glioma cells were previously transduced with lentivirus expressing eGFP and firefly luciferase, thereby allowing tumor burden to be monitored using *in vivo* luminescence imaging (IVIS Spectrum, PerkinElmer).

Intraventricular cannula (PlasticsOne, Torrington, CT) were placed at -0.6 mm posterior from bregma, 1.2 mm lateral to bregma and -2mm depth after tumors were established based on *in vivo* luminescence signaling^38^. Mice were then treated with weekly infusions of 2.5 million CD8 T-cells for a total of 4 weeks and monitored via weekly IVIS imaging.

For all *in vivo* experiments, a total of 6-9 mice per arm were utilized.

### Immunofluorescence

Mice were perfused with 4% PFA/PBS and brains were removed and post-fixed in 4% PFA/PBS overnight. Tissue was then sectioned on a vibratome at 50 µm thickness. Serial sections were stained overnight at 4C. Primary antibodies (anti-GFP antibody) were used according to manufacturer’s instructions.

Cells were plated on Matrigel and allowed to settle overnight, they were subsequently fixed in 4% PFA (Electron Microscopy Sciences) for 10 minutes and washed with PBS and PBS-T (3X 10 mins). Cells were blocked in 5% BSA and .1% Triton for 1 hour (Sigma-Aldrich), stained in primary antibody overnight at 4 degrees and washed with PBS-T (3x 10 mins). Finally, cells were incubated at room temperature for 1 hour in secondary antibody, washed in PBS-T, and mounted in ProLong Gold (Invitrogen).

### Statistical Analysis

All statistical analysis was performed using GraphPad Prism8 (GraphPad Software, La Jolla, CA) or Systat 13.0 (Systat software, Chicago, IL). We used “ploty” to generate the violin plots (https://chart-studio.plotly.com/create/#/). All *in vitro* assays were replicated a minimum of three times.

## Funding sources

DOD CDMRP Neurofibromatosis Program, Foerderer Award (CHOP), Matthew Larson Foundation, Rally Foundation, Kids Join the Fight Foundation, R25 NINDS, K12 NCI.

**Supplementary Video 1**: Mouse MCIPs migrate to 7316-1746 RFP positive tumor spheres in a 2D assay.

**Supplementary Video 2:** Mouse MCIPs migrate to 7316-1746 RFP positive tumor spheres in a spheroid assay.

**Supplementary Video 3**: Control CD19-BiTE expressing 293Ts co-cultured CD8 T-cells fail to eliminate glioblastoma 7316-1746 (red).

**Supplementary Video 4**: Targeting EGFR-BiTE expressing 293Ts co-cultured CD8 T-cells potently eliminate glioblastoma 7316-1746 (red).

## Supporting information

Video 1: Mouse 2D migration assay

Video 2: Mouse spheroid migration assay

Video 3: CD19 BiTE killing assay

Video 4: EGFR BiTE killing assay

